# Variation at the Klotho gene locus does not affect cognitive function in up to 335,074 British Caucasians in the UK Biobank

**DOI:** 10.1101/838409

**Authors:** Hasnat A Amin, Fotios Drenos, Alexandra I Blakemore

## Abstract

The proportion of older adults in Western populations is increasing and there is, therefore, a need to define factors affecting maintenance of physical and cognitive health in old age. Variations in the Klotho (*KL*) gene, and specifically the *KL*-VS haplotype, have been identified by several authors as potentially influencing cognitive function and decline. We have attempted to verify the reported associations between *KL* variants, including the *KL*-VS haplotype, and cognitive function in up to 335,074 British Caucasian participants aged 40-79 years from the UK Biobank. We do not find evidence that *KL*-VS affects cognitive function or its decline with increasing age. We examined a further 244 *KL* variants and found that rs117650866 was associated with Prospective Memory, but could not replicate this in follow-up samples. In conclusion, there is insufficient evidence in the UK Biobank to support the concept that *KL* variants affect cognitive function or its rate of decline.

## Introduction

The demographics of Western populations are changing, with an increase in the proportion of older adults. There is, thus, a need to define the factors affecting maintenance of physical and cognitive health in old age. Cognition can be defined as any process that is required for an individual to be aware of their situation and to use that information to respond to it (1). As individuals get older, memory, learning and processing speed decline (2); often leading to reduced independence and increased reliance on families and social care. As life expectancy increases, it becomes ever more necessary to explore some of the factors that might explain variation in cognitive function and cognitive decline in adults.

Several authors have highlighted variants in the Klotho (*KL*) gene as associated with ageing. *KL* is located on chromosome 13 in humans, and encodes a single-pass transmembrane protein that acts as an FGF23 co-receptor (3–5). It was first identified in mice by Kuro-o *et al.* (6) who showed that decreased *kl* expression resulted in a condition resembling premature ageing. In humans, *KL* variants have been reported to be associated with longevity, cardiovascular risk factors and cancer (7–10).

In addition, multiple studies have been carried out exploring the relationship between *KL* variants and cognitive function and decline, mostly focusing on the *KL*-VS haplotype, which refers to a pair of functional variants that result in F352V (rs9536314) and C370S (rs9527025) substitutions. Previous evidence has been varied: some authors have suggested that among adults aged 70 years or more, people homozygous for V (valine) at position 352 have poorer cognitive function (11,12), but also suggests that V352 heterozygotes have better cognitive function than those who are homozygous for (F) phenylalanine at position 352 (11,13). On the other hand, Mengel-From *et al.* (14) showed that, in Danish populations aged between 92-100 years, V352 heterozygotes had poorer cognition and Almeida *et al.* (15) showed that among men aged 71-87 years, V352 carriers were more likely to get dementia. De Vries *et al.* (16) showed that V352 heterozygotes have a slower rate of cognitive decline, but Porter *et al.* (17) did not find any such relationship in their data.

In addition to the *KL*-VS haplotype, there are reports of associations between variants in the *KL* promoter region and cognition. Mengel-From *et al.* (14) reported that carriers of the rs398655 C allele had better cognitive function than non-carriers and Hao *et al.* (18) reported that those with who are homozygous for the G (guanine) allele at G-395A (rs1207568) have an increased risk of cognitive impairment.

At present, conflicting reports indicate that, at a population level, the relationship between the *KL*-VS haplotype and cognitive function or cognitive decline is not particularly clear: there is, therefore, a need to explore this area further using significantly bigger sample sizes. Here, we aim to verify the reported associations between *KL* variants, including the *KL*-VS haplotype, and cognitive function in up to 335,074 UK Biobank (UKB) participants aged between 40 and 79 years, by carrying out a phenome scan of cognitive measures, including reaction time and various memory tests. We also aim to search for novel associations between the *KL* genetic variants and cognitive function using the same approach.

## Subjects and Methods

### Population and study design

This study was carried out using data from the UK Biobank (UKB). UKB is a large prospective cohort study that recruited ~502,600 UK residents aged between 40 and 69 years of age between 2006 and 2010. The participants provided blood, urine and saliva samples, and underwent various physical assessments, as well as touchscreen questionnaires and verbal interviews (19).

### Phenotypes

Table 1 summarises the phenotypes relating to cognitive function (referred to as cognitive measures) that were used for our analyses. For some cognitive measures, a baseline measurement was carried out (referred to as ‘Baseline’) at one of 22 assessment centres as well as 2 follow-up measurements (referred to as ‘Follow-Up 1’ and ‘Follow-Up 2’) for a subset of participants. For Fluid Intelligence, Pairs Matching and Numeric Memory, an online assessment was performed in addition to the measurements undertaken at the assessment centres. For Pairs Matching, there were 3 rounds; the first round had 3 pairs that the participants needed to match and the second and third rounds had 6. For Trail Making, only data from the online measurement was available to us. We did not include participants in the analysis for a given cognitive function test if they abandoned the test and/or if they completed the test with a pause. Each round/follow-up of each measure was treated as a separate phenotype unless otherwise stated.

### Genotyping and quality control

488,377 individuals were genotyped for up to 812,428 variants using DNA extracted from blood samples on either the UK Biobank Axiom array (438,427 participants) and the UK BiLEVE Axiom array (49,950 participants). Variant quality control metrics were provided by UKB as described previously (20). For genotyped variants, all variants that did not pass standard quality control checks carried out by Affymetrix and the Wellcome Trust Centre for Human Genetics were excluded. Specifically, hypothesis testing was carried out to check for differences in genotyping due to batch effects, plate effects, sex effects and array effects as well as any departures from Hardy-Weinberg Equilibrium using a p-value threshold of 10^−12^. In addition, variants with a missingness of >1% and/or a minor allele frequency of <0.01 were also excluded. For imputed variants, all variants with an INFO score of <0.8 were excluded. The *KL* gene is located at 13:33590571-33640282 (GRCh37.p13) and 246 variants passed QC within ±5 Kb of *KL*. These were selected for the association analyses.

Sample quality control metrics were provided by UKB and were generated as described by Bycroft *et al.* (20). Samples were excluded from the analysis if they were determined to be outliers for missingness and/or heterozygosity and/or if they had any sex chromosome aneuploidies as well as if the genetically inferred sex differed from the reported sex. Samples which did not have a genetically-determined White British ancestry were also excluded. A list of related individuals was also provided by UK Biobank and one individual from each related pair was excluded at random.

### Statistical Analyses

PLINK 2.0 was used to fit an additive linear model between the cognitive measures and the genotypes in all individuals. This was then repeated for a subset of individuals who were aged 69 years or more at the time of performing the cognitive test. Unless otherwise specified, all association analyses (i.e. additive linear models) were adjusted for the first 4 genetic principal components (PCs) (UKB Field 22009) and the genotyping chip on which the participant was genotyped on. The cognitive measures and any quantitative covariates were standardised to a mean of 0 and a variance of 1 before any linear modelling was performed.

Since multiple testing was undertaken, we applied statistical correction for this. A principal component (PC) analysis showed that 24 PCs represented >90% of variation in the 28 cognitive measures. To determine the number of independent variants, all pairs of variants within the locus with R^2^ > 0.1 were listed, one variant from each pair was removed and this process was repeated until there were no pairs of variants remaining. When this is implemented using --indep-pairwise 60 kb 1 0.1 in PLINK 2.0, 15 independent variants remain. A p-value threshold of 0.05 is used and Bonferroni-corrected when necessary for the appropriate number of independent tests in each case (up to 360 independent tests: 15 independent variants and 24 PCs). Supplementary Figure 1 summarises the analyses and the threshold used for each of them.

**Figure 1.**
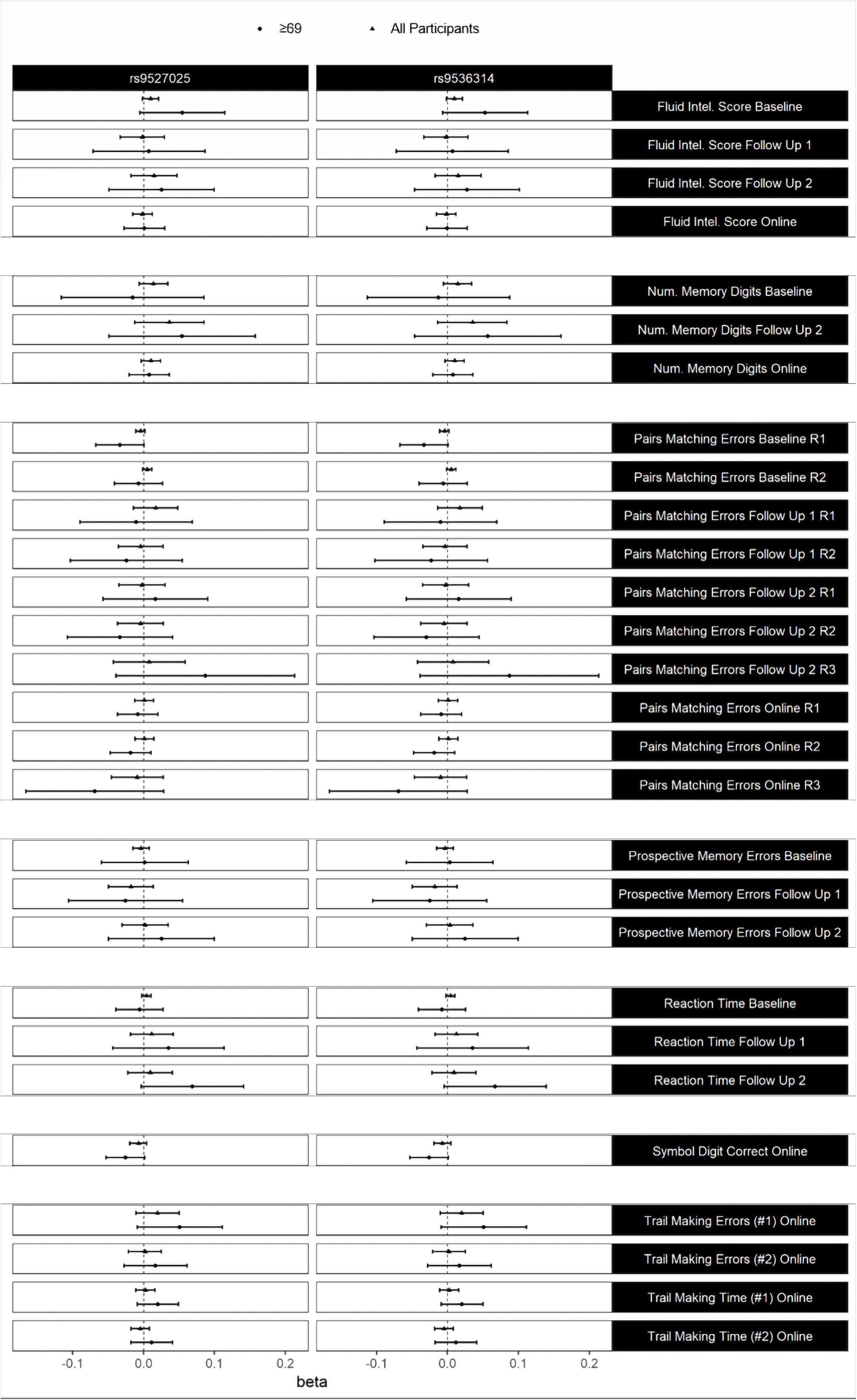
Standardised beta coefficients with 95% Confidence Intervals when regressing cognitive measures on rs9536314 and on rs9527025 in the UK Biobank with (All Participants) and without (≥69) including participants less than 69 years old and adjusted for age, age^2^ and sex.

## Results

After QC, there were 335,074 individuals remaining for analysis. A summary of the sample by phenotype is provided in Table 2.

Since the 2 variants making up the *KL*-VS haplotype are well-characterised functional *KL* variants in humans, we investigated whether either of them was associated with any of the cognitive function measures available. Neither rs9536314 nor rs9527025 were significantly associated at a p-value threshold of 0.05/24 with any of the cognitive measures when unadjusted (Supplementary Table 1) and when adjusted for age, age^2^ and sex (Figure 1).

Previous studies have largely concentrated on older individuals: to test whether these variants exerted their effects only in later life, we repeated the analyses, but only included individuals who were aged ≥69 years at the time that they performed the cognitive test. However, we again found that neither rs9536314 nor rs9527025 were significantly associated at a p-value threshold of 0.05/24 with any of the cognitive measures available (Figure 1 & Supplementary Table 1). It was not possible to increase this age threshold beyond 69 years because all participants were aged between 40 and 69 years at the time of recruitment.

Although the associations were not statistically significant, the effect size appeared to increase when excluding individuals under the age of 69 years. We therefore repeated the analyses but included a genotype*age interaction term to test whether the effect of *KL*-VS variants on the cognitive function measures available changed with age. We found that age does not have a statistically significant effect on the relationship between *KL*-VS and any of the cognitive function measures available at a p-value threshold of 0.05/24 adjusting for age, age^2^ and sex (Supplementary Table 2).

We next sought to test whether rs9536314 or rs9527025 were associated with change in any of the cognitive measures over age. For all measures for which more than one data point was available per participant (i.e. participants had performed a given cognitive test on more than one occasion), a rate of change was calculated for each participant (where the rate of change is the change in the cognitive measure, M2-M1, divided by the age difference, T2-T1, in years: on average, the difference between two measurements is 8.3 years). We found that neither rs9536314 nor rs9527025 was significantly associated at a p-value threshold of 0.05 with a change in any of the available cognitive measures over age, either when unadjusted or when adjusted for age_T1_, age_T1_^2^ and sex (Supplementary Table 3).

We next tested to see if any other *KL* variants were statistically associated at a p-value threshold of 0.05/360 with any of the available cognitive measures. We found that rs117650866 was associated with Prospective Memory (UKB Field 4291) adjusting for age, age^2^ and sex (referred to as Discovery in Table 3). There were no other significant associations, with or without adjustment for age, age^2^ and sex (Supplementary Table 4 & 5); there were also no significant associations when excluding individuals under the age of 69 years (Supplementary Table 4 & 5).

We then attempted to internally replicate the rs117650866 association. To do this, we repeated the association analysis in participants who performed the Prospective Memory test only on one occasion (i.e. those participants who performed the test at either Baseline only, or at Follow Up 1 only, or at Follow Up 2 only). The rs117650866 association was present in those tested at Baseline only (beta = 0.112, s.e. = 0.0229, power = 0.92, v = 99054), but was absent (p > 0.05) in those tested only at either Follow Up 1 only (beta = −0.059, s.e. = 0.0794, power = 0.12, v = 8179) or at Follow Up 2 only (beta = −0.117, s.e. = 0.101, power = 0.09, v = 5656) (Table 3). Despite the lack of power to detect the association observed at Baseline, the inconsistent direction of the effect between Baseline and Follow Up 1 and 2 suggest that the association observed at Baseline was likely to be a false positive. The power calculations were carried out using the pwr.f2.test function from the pwr package in R with u set to 9, f2 set to 0.000216 and sig.level set to 0.05.

We next sought to test whether any *KL* variants were significantly associated at a p-value threshold of 0.05/15 with a change in any of the cognitive measures (for which there are repeat measurements) over age, in the same way that *KL*-VS was tested. We did not find any significant associations (Supplementary Table 6).

## Discussion

Previous evidence suggested that *KL*-VS and other *KL* variants are associated with cognitive function during the later stages of life. Our aim was to explore these findings in a younger and much larger cohort, namely the UK Biobank. We did not find evidence of a relationship between *KL*-VS and cognitive function, nor did we find any evidence that the age of an individual had a significant effect on this relationship. The association we found between Prospective Memory and rs117650866 did not replicate consistently, nor is there any evidence of it in previously published studies, so it is likely to be a false positive. We also did not find evidence of any other *KL* variants being associated with cognitive function nor with cognitive decline.

An important point is that previous studies which have identified relationships between *KL* variants and cognition use populations that are much older (usually aged 70 years or more), whereas the population we examined is relatively young (the larger Baseline samples had a mean age of 57 years). We attempted to address this limitation by repeating our analyses, but only including individuals over the age of 69 years; we still did not find associations probably because only about 1% of this subset in the Online tests are over 75 years of age and no individuals are over 79 years of age. Indeed, whenever authors report an absence of statistically significant associations between *KL* variants and cognition, the mean age of the cohorts that they analyse are closer to the one we analysed. For example, Deary *et al.* (12) examined 2 cohorts and the cohort who undertook cognitive testing at age of 64 years did not show statistically significant associations between *KL*-VS and cognition. Dubal *et al*. (13) also did not find an association in one of the 3 cohorts that they analysed, and the mean age of this cohort was 63 years.

Deary *et al.* (12) provided evidence suggesting that *KL*-VS may influence cognitive decline. We did not find any evidence to support this. This may be because the difference between the repeated measurements available to us was about 8 years whereas Deary *et al*. compared the cognitive abilities of individuals first tested when aged 11 years and then at the age of 79 years. It is also important to note that whilst some authors do report relationships between *KL*-VS and cognitive decline (12,16), other authors do not find any such relationship (14,17).

The UKB dataset, despite the advantage of its size, does have biases. In particular, the participants are generally healthier than average (21). There is evidence to suggest that the effect of *KL* variants on cognitive function/decline may be as a result of affecting the severity of a pre-existing psychopathology (22,23) and individuals suffering from early dementia, etc. would be either unlikely or even unable to volunteer to participate.

In conclusion, there is insufficient evidence in the UK Biobank to support the concept that *KL* variants affect cognitive function or its rate of decline in British Caucasian individuals aged between 40 and 79 years. Further follow-up testing would be required to verify the reported effects of *KL* on cognitive function and decline that are reported in very elderly individuals.

## Acknowledgements

This study was carried out under UK Biobank application 19968 and we would like to thank both the UKB participants and the UKB staff. The application was paid for by Calico LLC (South San Francisco, California, United States). Hasnat Amin is the recipient of a PhD studentship from the College of Health and Life Sciences, Brunel University London.

---

<u>Supplementary Figure 1</u>

A breakdown of the analyses carried out in this study. LD = linkage disequilibrium. PCs = principal components.

## Tables

<u>Table 1</u>

A description of the cognitive measures from the UK Biobank used in this study.

<u>Table 2</u>

Demographics of the cohort by phenotype. The distribution of the phenotype is presented in the original units.

<u>Table 3</u>

Standardised coefficients (beta), standard errors (s.e.), sample sizes (n) and p-values (p) when regressing cognitive measures on rs117650866 in the UK Biobank adjusted for age, age^2^ and sex in the full baseline sample (Discovery), and in 3 independent samples (i.e. in participants who were present at baseline only (Baseline) and in participants who were present at the first follow up only (Follow Up 1) and in participants who were present at the second follow up only (Follow Up 2)).

<u>Supplementary Table 1</u>

Standardised coefficients (beta), standard errors (s.e.), sample sizes (n) and p-values (p) when regressing cognitive measures on rs9536314 and on rs9527025 in the UK Biobank with (All Participants) and without (≥69 years old) including participants less than 69 years old and not adjusted.

<u>Supplementary Table 2</u>

Standardised coefficients (beta), standard errors (s.e.), sample sizes (n) and p-values (p) when regressing cognitive measures on rs9536314 and on rs9527025 in the UK Biobank with a genotype*age interaction term and adjusted for age, age^2^ and sex.

<u>Supplementary Table 3</u>

Standardised coefficients (beta), standard errors (s.e.), sample sizes (n) and p-values (p) when regressing the rate of decline of cognitive measures on rs9536314 and on rs9527025 in the UK Biobank with (Adjusted) and without (Unadjusted) adjusting for age_T1_, age_T1_^2^ and sex.

<u>Supplementary Table 4</u>

Standardised coefficients (beta), standard errors (s.e.), sample sizes (n) and p-values (p) when regressing cognitive measures on 246 *KL* variants in the UK Biobank with (All Participants) and without (≥69 years old) including participants less than 69 years old and not adjusted.

<u>Supplementary Table 5</u>

Standardised coefficients (beta), standard errors (s.e.), sample sizes (n) and p-values (p) when regressing cognitive measures on 246 *KL* variants in the UK Biobank with (All Participants) and without (≥69 years old) including participants less than 69 years old and adjusted for age, age^2^ and sex.

<u>Supplementary Table 6</u>

Standardised coefficients (beta), standard errors (s.e.), sample sizes (n) and p-values (p) when regressing the rate of decline of cognitive measures on 246 *KL* in the UK Biobank with (Adjusted) and without (Unadjusted) adjusting for age_T1_, age_T1_^2^ and sex.

