## Supplementary figures and images for "Variation at the Klotho gene locus does not affect cognitive function in up to 335,074 British Caucasians in the UK Biobank"

### Supplementary Figure 1

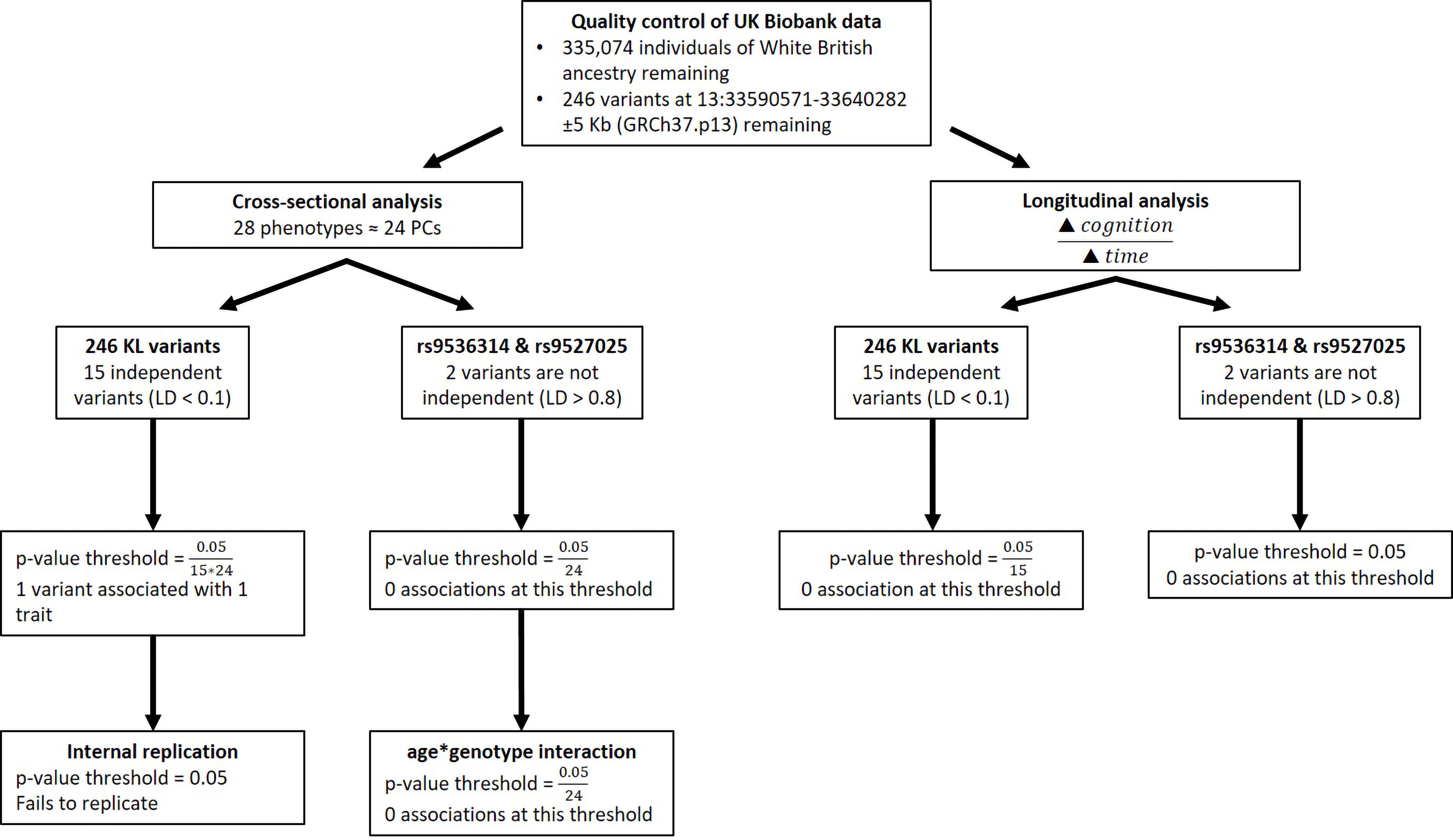
